# A very fast time scale of human motor adaptation: within movement adjustments of internal representations during reaching

**DOI:** 10.1101/269134

**Authors:** F. Crevecoeur, J.-L. Thonnard, P. Lefèvre

## Abstract

Humans and other animals adapt motor commands to predictable disturbances within tens of trials in laboratory conditions. A central question is how does the nervous system adapt to disturbances in natural conditions when exactly the same movements cannot be practiced several times. Because motor commands and sensory feedback together carry continuous information about limb dynamics, we hypothesized that the nervous system could adapt to unexpected disturbances online. We tested this hypothesis in two reaching experiments during which velocity-dependent force fields were randomly applied. We found that within-movement feedback corrections gradually improved, despite the fact that the perturbations were unexpected. Moreover, when participants were instructed to stop at a via-point, the application of a force field prior to the via-point induced mirror-image after-effects after the via-point, consistent with within-trial adaptation to the unexpected dynamics. These findings suggest a fast time-scale of motor learning, which complements feedback control and supports adaptation of an ongoing movement.

**Significance Statement:** An important function of the nervous system is to adapt motor commands in anticipation of predictable disturbances, which supports motor learning when we move in novel environments such as force fields. Here we show that movement control when exposed to unpredictable disturbances exhibit similar traits: motor corrections become tuned to the force field, and they evoke after effects within an ongoing sequence of movements. We propose and discuss the framework of adaptive control to explain these results: a real-time learning algorithm, which complements feedback control in the presence of model errors. This candidate model potentially links movement control and trial-by-trial adaptation of motor commands.

## Introduction

Neural plasticity in the sensorimotor system enables adaptive internal representations of movement dynamics and acquisition of motor skills with practice. In the context of reaching movements, studies have documented that healthy humans and animals can learn to anticipate the impact of a force field applied to the limb, and recover straight reach paths within tens to hundreds of trials (Lackner and DiZio, 1994; Shadmehr and Mussa-Ivaldi, 1994; Krakauer et al., 1999; Thoroughman and Shadmehr, 2000; Singh and Scott, 2003; Wagner and Smith, 2008). Importantly, studies on motor learning have consistently highlighted the presence of an after-effect, which mirrors the initial movement deviation and indicates that adaptation was supported by a novel internal model of movement dynamics (Lackner and DiZio, 2005; Shadmehr et al., 2010; Wolpert et al., 2011).

To date, motor learning and adaptation have been exclusively studied on a trial-by-trial basis, highlighting context-dependent learning rates and memory dynamics across movements (Smith et al., 2006; Kording et al., 2007; Diedrichsen et al., 2010; Wei and Koerding, 2010; Gonzalez Castro et al., 2014). It is clear that there exist medium to long time scale in the acquisition of motor skills and in the adaptation of motor commands to changes in the environment which impact motor performances across movements (Krakauer and Shadmehr, 2006; Dayan and Cohen, 2011). But, although this approach has revealed fundamental properties of sensorimotor systems, it has left unexplored the problem of online control during early exposure to the altered environment. More precisely, it remains unknown how the nervous system controls reaching movements in the presence of unexpected dynamics, or errors in the internal models, as in the early stage of motor adaptation.

It is often assumed that unexpected disturbances during movements are automatically countered by the limb’s mechanical impedance and by reflexes (Shadmehr and Mussa-Ivaldi, 1994; Burdet et al., 2000; Franklin and Wolpert, 2011; Milner and Franklin, 2005). However, recent work suggests that muscles viscoelastic properties, as well as the gain of the spinal reflex (latency ∼20ms for upper limb muscles) are low at spontaneous levels of muscle activation (Crevecoeur and Scott, 2014). Furthermore, long-latency reflexes (latency ≥50ms) are also based on internal models of the limb and environmental dynamics (Kurtzer et al., 2008; Cluff and Scott, 2013; Crevecoeur and Scott, 2013), even when disturbances are very small (Crevecoeur et al., 2012). Thus, the presence of model errors is equally challenging for rapid feedback control as it is for movement planning, and yet healthy humans can handle unexpected disturbances relatively well.

Thus, the outstanding question is whether and how the nervous system controls movements online when exposed to unexpected dynamics. In theory, an approximate internal model can be deduced during movement because motor commands and sensory feedback together carry information about the underlying dynamics. This problem was studied in the framework of *adaptive control* (Bitmead et al., 1990; Ioannou and J, 1996; Fortney and Tweed, 2012). The idea is to adapt the model continuously based on sensory feedback about the unexpected movement error. It is important to underline that adaptive control aims at correcting for the presence of model errors or fixed biases in the system model. When there is no such error, and when disturbances follow a known distribution, this framework reduces to standard (extended) stochastic optimal control (Todorov and Jordan, 2002; Todorov, 2005; Izawa et al., 2008).

In light of the theory of adaptive control and considering the very rich repertoire of rapid sensorimotor loops (Scott, 2016), we explored the possibility that mechanical disturbances to the limb update the internal representations of the ongoing movement. An online estimation of task-related parameters was previously reported in the context of virtual visuomotor rotations (Braun et al., 2009), where participants had to estimate the mapping between hand and cursor motion. Here we address this problem in the context of reaching in a force field to probe specifically the internal representations of limb dynamics. To investigate this question, we used a random adaptation paradigm and addressed whether participants’ responses to unpredictable disturbances reflected the presence of adaptation. We show that participants learned to produce feedback responses tuned to the force field, and illustrate how their behaviour could be explained in the framework of adaptive control. We highlight the limitations of this and alternative candidate models, and underline the associated computational challenges for the nervous system that have yet to be fully characterized.

## Methods

### Experimental Procedures

A total of 36 healthy volunteers (15 females, between 23 and 36 years) provided written informed consent following procedures approved by the ethics committee at the host institution (University of Louvain). Participants were divided into two groups of 14 for the two main experiments, and one group of 8 for the control experiment. The three experiments are variants of the same task. Participants grasped the handle of a robotic device (KINARM, BKIN Technologies, Kingston, Canada) and were instructed to perform reaching movements towards a visual target. The movement onset was cued by filling in the target (Fig. 1a), and feedback was provided about movement timing: good trials were between 600ms and 800ms between the go signal and the moment when they entered the target. Direct vision of the hand was blocked, but the hand-aligned cursor was always visible.

**Figure 1:**
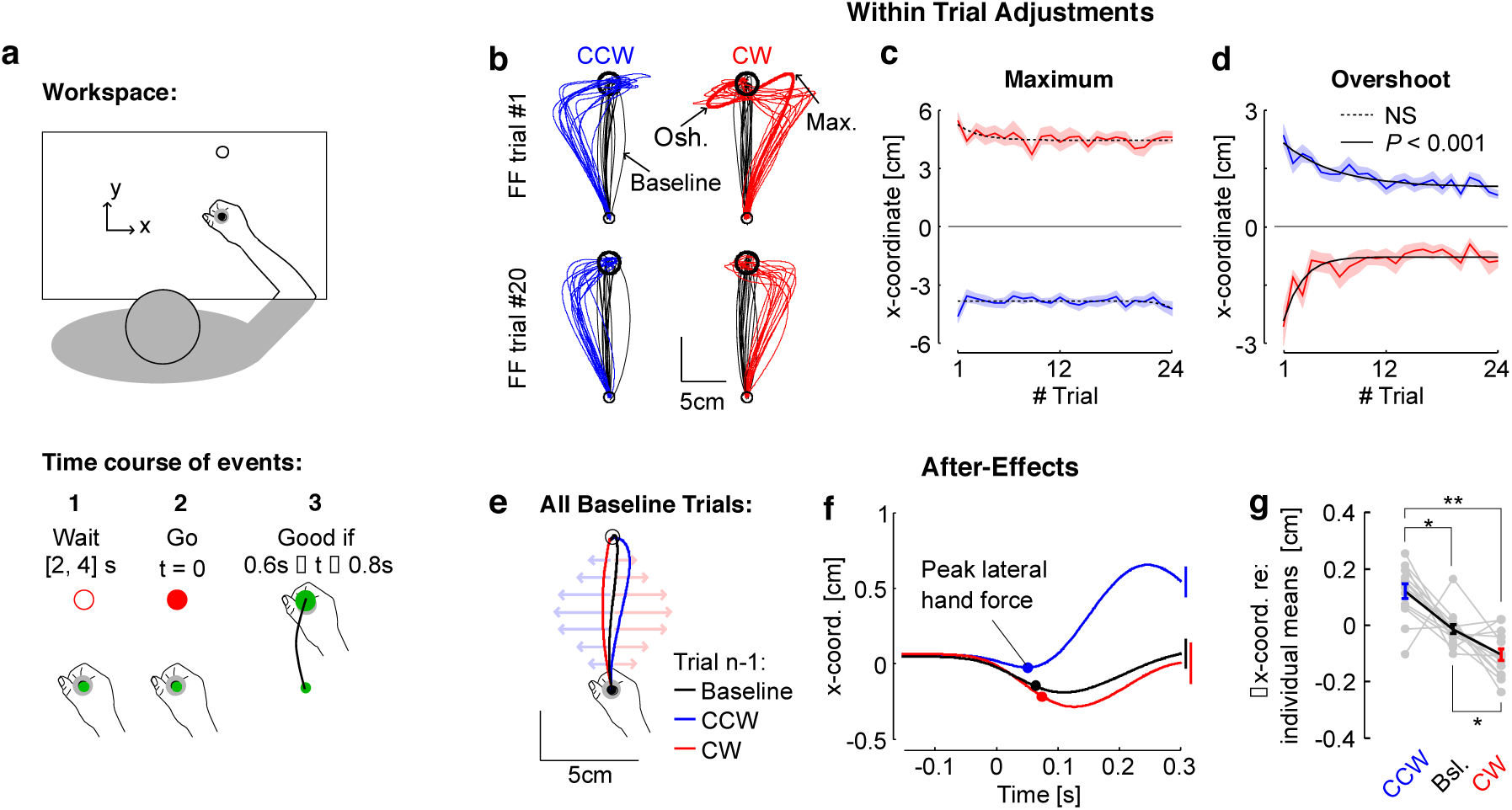
Experiment 1, Behavior. a: Illustration of a successful trial in Experiment 1 (n=14). Participants were instructed to wait for the go signal, and then to reach for the goal within 0.6 s to 0.8 s of the go signal. Feedback about movement timing was provided to encourage participants to adjust their movement speed, but all trials were included in the dataset. b: Hand paths from all participants for the first force field trial (top) and the force field trial number 20 (bottom), were selected to illustrate the change in online control. Counter-clockwise (CCW) and clock-wise (CW) FF trials are depicted in blue and red, respectively. The black traces (baseline) are randomly selected trials from each participant. Maximum deviation in the direction of the force field (Max.) and the maximum target overshoot (Osh.) are illustrated. c: Maximum hand path deviation for CCW and CW force fields across trials. The dashed traces illustrate the non-significant exponential fits. d: Maximum target overshoot across FF trials. Solid traces indicate significant exponential decay. Shaded blue and red areas around the means indicate the SEM across participants. e: Illustration of the after effect. Baseline trials were categorized dependent on whether they were preceded by CCW or CW force fields, or by at least 3 baseline trials. f: Continuous traces of x-coordinate as a function of time and extraction at the moment of peak lateral hand force (dots). Error bars at 300ms illustrate the SEM across participants. g: individual averages of lateral hand coordinate (gray), and group averages (mean±SEM), after subtracting each individual’s grand mean for illustration). One (two) star(s) indicate significant difference at the level *P* < 0.05 (0.01) based on paired t-test, adjusted for multiple comparison with Bonferroni correction.

#### Experiment 1

The goal of this experiment was to characterize online control in the presence of unexpected changes in reach dynamics. Participants (n = 14) performed baseline trials randomly interleaved with orthogonal force field trials mapping the forward hand velocity onto a lateral force 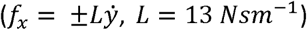. Movements consisted of 15cm reaches in the forward direction. To make the task less predictable, other trial types were also randomly interleaved including trials with via-points, and trials with a constant background load applied to the hand. The via-point trials of Experiment 1 were located on the either side of the reach path (coordinates in cm: [±4, 12]), and participants were simply instructed to go through them. For the trials with background force, there was a 500ms build up prior to the onset cue, and the switch off was after trial end (*f*_*x*_ = ±4*N*). These trials occurred with the same frequency as the force field trials. Participants performed six blocks of 80 trials including 56 baseline trials and 4 force field trials per direction of the force field and per block, summing to a total of 24 force field trials per direction (clockwise, CW, and counterclockwise, CCW) and participant. For this experiment, feedback was provided about success to encourage a consistent velocity across trials but all force field trials were included in the analyses.

#### Experiment 2

This experiment was designed to address whether the improvement in online correction observed in the first experiment could evoke a near instantaneous after effect. Movements consisted of 16cm reaches in the forward direction, and a via-point (radius: 1cm) was located at 10cm on the straight line joining the start and goal targets. Participants (n=14) performed six blocks of 80 trials composed of 45 baseline trials, 5 force field trials, 5 baseline trials with a via-point, 5 force field trials with a via-point, and 5 force field trials with a via-point and with a switching off of the force field after the via-point. All trials and force field directions were randomly interleaved. Participants were given scores for good trials (1 point), i.e. when they reached the goal within the prescribed time window (maximum 1.2s for trials with a via-point), and bonuses when they stopped successfully at the via-point (3 points). The bonus was awarded if the hand speed inside the via-point dropped below 3cm/s. The experimental setup monitored the hand speed in real time allowing to turn off the force field if (1) the hand cursor was in the via-point and (2) the hand speed dropped below 3cm/s while in the via-point (determined based on pilot testing). Feedback about a successful stopping at the via-point was given online. In this experiment, we included in the analyses all force field trials without via-point similar to Experiment 1. For the via-point trials with unexpected switch-off of the force field, we had to include only the trials for which participants successfully stopped according to the speed threshold of 3cm/s, since these were the only trials for which the force field was effectively turned off. This condition made the via-point trials quite difficult and we recorded an average of 21, 25 and 17 successful trials for counterclockwise, baseline and clockwise via-point trials, respectively (range across subjects: [5, 30], [18, 29], [2, 30]).

#### Control Experiment

This control experiment was designed to characterize participants’ behavior in a standard trial-by-trial learning paradigm. Participants (n = 8) performed a series of 180 force field trials (CW or CCW), followed by a series of 20 baseline trials to wash out learning, and finally they performed another series of 180 force field trials in the opposite direction as the first series. The trial protocol with target location, random wait time and prescribed time window to reach for the goal target was identical as that of Experiment 1. The order in which the series of CW or CCW trials were performed was counterbalanced across participants.

### Analysis and Statistical Design

Position and force signals were sampled at 1kHz. Velocity signals were derived from numerical differentiation (2^nd^ order centered finite difference algorithm). The presence of significant main effects was assessed with repeated-measures ANOVAs performed on individuals’ mean value of the variable of interest (rmANOVA). Post-hoc tests were performed based on paired t-test with Bonferroni correction for multiple comparisons. In experiment 1, one participant missed a force-field trial such that the analyses were performed on 23 trials for the corresponding force field direction. We assessed the evolution of several parameters such as maximum displacement in the lateral direction, maximum lateral target overshoot, and peak end forces, and path length by means of exponential fits. The exponential models were fitted in the least-square sense and the significance of the fit was determined based on whether the 99.5% confidence interval of the exponent responsible for the curvature of the fit included or not the value of zero (*P*<0.005). The non-significant fits were associated with *P*>0.05. An *R*^2^ statistics was derived as follows:

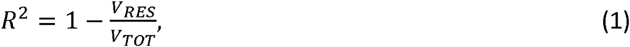

where *V*_*RES*_ denotes the variance of the residuals and *V*_*TOT*_ is the total variance of the data. We adopted Cohen’s *d* to quantify the effect size for paired comparison as the mean difference between two populations divided by the standard deviation of the paired difference between the populations (Lakens, 2013).

Adaptation to the force field disturbances was assessed based on the correlation between the commanded force and the measured force. Indeed we can decompose the forces acting on the handle as follows: the force induced by the robot dynamics (such as inertial force field and friction) called *f*_*R*_, the commanded force of the force field environment called *f*_*ENV*_, and the force at the interface between the hand and the robotic handle called *f*_*H*_, which is measured with the force transducer. At this stage there is no distinction of passive or active components of the force applied by the hand to the handle. The handle acceleration is equal to the sum of the forces acting on it:

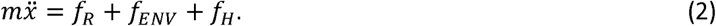

Then, the use of the correlation between *f*_*H*_ and *f*_*ENV*_ as an index of adaptation (with appropriated sign) can be justified by the observation that the differences between these two forces was well correlated with the lateral acceleration. Indeed, the commanded force in the environment was calculated offline as the forward velocity multiplied by the scaling factor that defines the force field (*L* = ±13). Then, we calculated the difference between the measured and commanded forces, and observed correlations with the lateral acceleration characterized by a mean R^2^ of 0.92 (range: [0.81, 0.95]), and a mean slope of 0.93±0.03 (mean±SD, averaged across direction). Note that this error is also in part induced by real-time sampling of velocity in the system. Thus the error made by ignoring the unmodelled forces induced by the KINARM (*f*_*R*_ in Eqn. 2) and sampling inaccuracies represented <10% of unexplained variance on average, and the 90% of explained variance was accounted for by the lateral acceleration. Since *f*_*R*_ and *f*_*ENV*_ must be the same for movements with similar kinematics, any change the correlation between the measured and commanded forces reflect changes in control (*f*_*H*_). We also used the data from the control experiment to validate this approach empirically.

The data of experiment 2 was also analyzed in more detail based on individual trials. The analyses of individual trials within each participant were based on Wilcoxon rank-sum test. We used mixed linear models of lateral hand velocity as a function of the lateral hand velocity prior to the via-point, and also as a function of the trial number based on standard techniques (Laird and Ware, 1982). The possibility that changes in internal representations impacted the very first trials was assessed by regressing velocity against the trial index. For this analysis, we changed the sign of the lateral velocity for clockwise force field trials such that for all trials, a positive modulation reflected the presence of an after effect. The dependent variable was the lateral hand velocity measured at the moment of the second velocity peak, the fixed effect was the trial index, and the model contained a random offset per to account for idiosyncrasy.

### Model

The model describes the translation of a point-mass similar to the mass of an arm (*m* = 2.5 kg) in the horizontal plane. The coordinates corresponded to the workspace depicted in Fig. 1, such that the reaching direction was along the *y* dimension. Dot(s) correspond to time derivative. The variables *f*_*x*_ and *f*_*y*_ are the forces applied by the controlled actuator to the mass, and these forces are a first order response to the control vector denoted by the variables *u*_*x*_ and *u*_*y*_. The state-space representation was thus as follows:

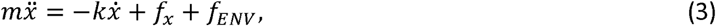

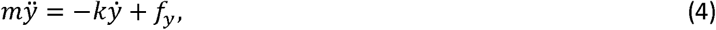

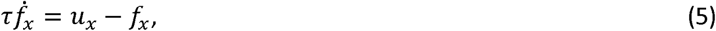

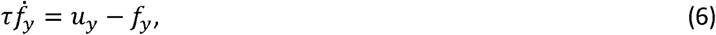

where *k* is a dissipative constant (0.1 Nsm^-1^), Equations 5 and 6 capture the first order muscle dynamics with time constant set to 0.1s, and *f*_*ENV*_ represents the unknown environmental perturbation: *f*_*ENV*_ = 0 for the null field, or 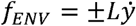 for force field trials.

Defining the state vector as 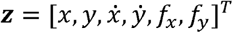, and the control vector ***u***= [*u*_*x*_, *u*_*y*_]^*T*^, we have ***ż*** = *A*_0_***z*** + *B*_0_***u***, with:

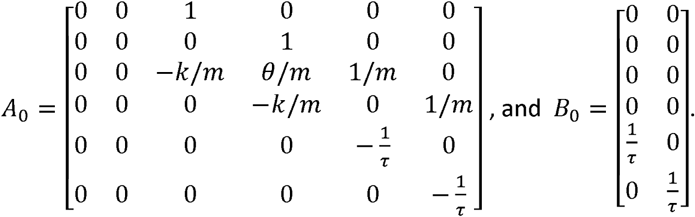

The parameter *θ* is either 0 for baseline (unperturbed) trials, or ±*L* for force field trials, in agreement with the definition of *f*_*ENV*_. The system was transformed into a discrete time representation by using a first order Taylor expansion over one time step of *δt*: *A*= *I* + *δtA*_0_, and *B* = *δtB*_0_ (*I* is the identity matrix). We used a discretization step of *δt* = 0.01*s*. The value of *θ* is unknown at the beginning of each trial. Thus we assume that it is unknown for the controller and the model error, Δ*A*, comes from a possible mismatch between expected and true values of *θ*.

The state vector and state space representation matrices were augmented with the coordinates of the goal (denoted by *x*\* and *y*\*), and the system was then re-written as follows:

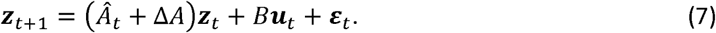

The subscript *t* is the time step, where *Â*_*t*_ is the current (time varying) expected dynamics, Δ*A* is the unknown model error containing the unmodeled environmental disturbance (*f*_*ENV*_), and ***ε***_*t*_ is a Gaussian disturbance with zero-mean and covariance Σ_*ε*_ := *BB*^*T*^. Recall that the true dynamics is the expected dynamics plus the model error, in other words we have *A* := *Â*_*t*_ + Δ*A*. As a consequence the model error also evolves across time as the adaptive controller changes the values of *Â*_*t*_.

We assumed for simplicity that the controller has perfect state measurement, meaning that *z*_*t*_ is known (fully observable case). The problem is that the controller does not know Δ*A* and must control the trajectory of the system in the presence of a model error. This problem was considered by Izawa and colleagues in the context of motor adaptation to random force fields (Izawa et al., 2008). These authors expressed that Δ*A* followed a known (Gaussian) distribution, which induced state-dependent noise and could be handled in the framework of extended stochastic optimal control (Todorov, 2005). Observe that this assumption implies that the force field intensity varied randomly across time, which was not consistent with our experiment for which the force field varied randomly across trials, but remained fixed within trials. In other words, stochastic optimal control is not consistent with our random adaptation design because Δ*A* depends on *f*_*ENV*_, which is not a stochastic variable but instead fixed bias or error in the model within each force field trial. Thus, for the present study, we expressed in the model the fact that the initial controller was derived assuming that the system dynamics corresponds to *A*, but that there was a potential error in the model. To further simplify the problem, we assume that the controller knows which parameter is unknown (*L*). Let *Â*_*t*_ denote the expected dynamics at time *t* dependent on the current estimate of the force field intensity noted 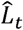, a Linear-Quadratic-Gaussian controller can be derived, giving control signals based *Â*:

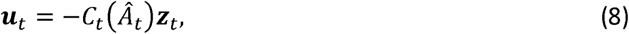

where −*C*_*t*_ (*Â*_*t*_)is the time series of feedback gains that, when applied to the state vector, defines the optimal control policy (Todorov, 2005). Our choice to simulate reaching movements in the context of optimal feedback control was motivated by previous work showing that this model accounted for a broad range of features expressed in human reaching movements such as time varying control gains, selective corrections, and flexibility (Diedrichsen, 2007; Liu and Todorov, 2007; Nashed et al., 2012).

Now, under this assumption, we may predict the next state vector, which we designate by 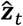. When the true dynamics is distinct from the expected dynamics (Δ*A* = *A* − *Â*_*t*_ ≠ 0), there is a prediction error, 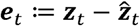, which can be used to correct the estimate of the model. We follow standard Least Square identification techniques (LS), and use the following rule to update the estimate of based on error feedback (Bitmead et al., 1990):

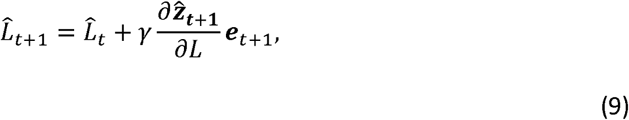

where *γ* is the online learning rate. This new estimation of the force field is then used to update *Â*_*t*+1_, and a new series of feedback gains is applied to the next state as control law: ***u***_*t*+1_ = −*C*_*t*+1_ (*Â*_*t*+1_)***z***_*t*+1_. In all, the closed loop controller consists in iteratively deriving optimal control gains based on an adaptive representation of the unknown parameters estimated online using Eqn. 9. It can be shown in theory that under reasonable assumptions, this adaptive optimal control scheme robustly stabilizes the closed loop system (Bitmead et al., 1990). For the simulations presented in this paper we verified that the time series of 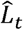 evolved in the direction of the true value during each movement.

The free parameters of the model are the cost-function used to derive the LQG controller, and the online learning rate *γ*. We fixed the cost-function to generate smooth movements with bell-shape velocity profiles, qualitatively comparable with human movements, and allowed *γ* to vary in order to capture the greater online adjustments observed across trials in Experiment 1.

We use the following cost function to simulate reaching movements without via-point:

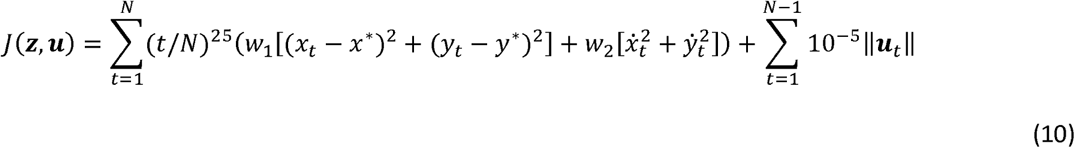

with *N* = 61, consistent with the average movement duration from Experiment 1 (∼600ms). The parameters *w*_1_ and *w*_2_. were set to 1000 and 20 to reproduce smooth movements with bell-shape velocity profiles. The first term produces a polynomial buildup of the terminal cost with value very close to 0 until about 80% of the terminal step, followed by a smooth buildup of the terminal cost for the last ∼20% of the reach time. For the via-point experiment, the cost function was identical except that we used *N* = 100 to account for larger duration of reaching with a via-point, and we added the penalty that the system had to be close to the via-point at *t* = 60 with the same cost as the terminal cost in Equation 9. There was no quantitative fitting of parameter values to the data.

The simulation of standard learning curves was performed by letting *Â*_*t*_ evolve during movements and storing the value obtained at the end of the trial for the beginning of the next trial (*γ* = 0.05). The reduction in peak end-force observed in Experiment 1 was simulated by using a range of values of *γ* to show that the range observed experimentally could be explained by this model (*γ* = 0.1, 0.25, and 0.5). Importantly, the measured force at the handle and the force produced by the controller were compared based on the observation that ignoring the robot dynamics only induced small errors. Finally, Experiment 2 was simulated as a series of null field trials interleaved with force-field trials with switching-off at the via-point. Two values of *γ* were chosen to illustrate that the modulation of lateral velocity after the via-point increased with this parameter (*γ* = 0.2 and 0.5). The simulated baseline trials in the simulation of Experiment 2 were preceded by three baseline trials to induce washout in the model. We extracted the local minimum of lateral velocity and the lateral velocity at the moment of the second peak velocity to illustrate the after effect in the simulations.

We tested an alternative model that did not involve any learning, but in which control was adapted following a time-evolving cost-function. The parameter *γ* was set to zero (i.e. *Â*_*t*_ was fixed), and the cost parameters were increased or decreased at each time step prior to re-computing the control gains. More precisely, the state related cost parameters (*w*_1_ and *w*_2_., Eqn. 10) were multiplied by 0.95 or 1.05 at each time step, and the feedback gains were recomputed at each step to take the time-evolving cost-function into account.

## Results

### Experiment 1

Participants grasped the handle of a robotic device and performed forward reaching movements towards a visual target (Fig. 1a). In the first experiment, clockwise and counterclockwise orthogonal force fields (FF) were randomly applied as catch trials, such that neither the occurrence nor the direction of the force field could be anticipated. Reach paths were of course strongly impacted by the presence of the force field, however the online corrections became smoother with practice (compare trials #1 and #20 in Fig. 1b). To quantify this, we extracted the maximum hand deviation along the direction of the force field in the lateral direction, and the target overshoot also measured in the same direction near the end of the movement (Fig. 1b: Max. and Osh., respectively). We found that the maximum deviations as well as their timing did not evolve significantly across force field trials (Fig. 1c, *P*>0.05), whereas the overshoot displayed a significant exponential decay (Fig. 1d, *P*<0.001, *R*^2^ = 0.17 and 0.56 for clockwise and counterclockwise force fields, respectively). The absence of a clear change in the maximum deviation suggests that there was no clear anticipation of the disturbances, and that measurable changes in control became apparent later. Only small adjustments impacting the maximum lateral displacement could be observed, in particular for counterclockwise trials that exhibited a reduction of ∼1cm on average. However the evolution of this variable across trials did not follow the same exponential decay as the target overshoot.

We also observed that exposure to force field trials induced standard after-effects on the next trial, as previously reported in both standard and random adaptation experiments (Lackner and DiZio, 1994; Shadmehr and Mussa-Ivaldi, 1994; Scheidt et al., 2001). To observe this, we separated trials performed in the null field dependent on whether they were preceded by force field trials or by at least 3 null field trials (Fig 1e), and extracted the x-coordinate of the hand path at the moment of lateral peak hand force (Fig. 1f) (this moment was well defined as it was induced by the inertial anisotropy of the KINARM handle). The lateral coordinate was clearly impacted by the occurrence of a force field trial, and the effect was opposite to the perturbation encountered in the preceding trial consistent with standard after effects (Fig. 1g, rmANOVA: *F*_(2,26)_=18.3, *P*<10^−4^). It can be also observed that the evolution of maximum target overshoot was not symmetrical (Fig. 1d), and the unperturbed trials were slightly curved (Fig. 1f). The origin of this asymmetry is unclear. On the one hand the perturbations did not engage the same muscles and differences in biomechanics might induce directional biases. On the other hand, the KINARM has anisotropic mass distribution, which induces an inertial force field. In spite of these differences, all effects reported were qualitatively similar across perturbation directions.

Several control strategies could produce a reduction in target overshoot, including an increase in control gains. However, the analyses of the measured force at the handle indicated a different strategy. We observed a reduction in the interaction force at the handle near the end of the movement, which suggested a decrease in lateral force used to counter the force field. Indeed, there was no significant change in the peak forward hand velocity across trials for both clockwise (rmANOVA, *F*_(23,299)_ = 0.56, *P* = 0.9) and counterclockwise force fields (*F*_(22,286)_ = 0.87, *P* = 0.62). Thus the perturbation applied to the limb remained statistically similar. However, the absolute peak hand force near the end of the movement decreased significantly. This is illustrated in Fig. 3, which displays the average perturbation and measured forces as a function of time for the first and last force field trials (Fig. 2a and b). There was a significant exponential decay of the terminal peak hand force (Fig. 2c, *R*^2^ =0.17 and 0.11 for clockwise and counterclockwise perturbations). As a consequence, the correlation between the measured lateral force and the commanded force field perturbation significantly increased across trials (Fig. 2d). We show below with the control experiment that these observations also characterize motor adaptation in a standard context of trial-by-trial learning, and that the increase in correlation shown in Fig. 2d can be used as an index of adaptation.

**Figure 2.**
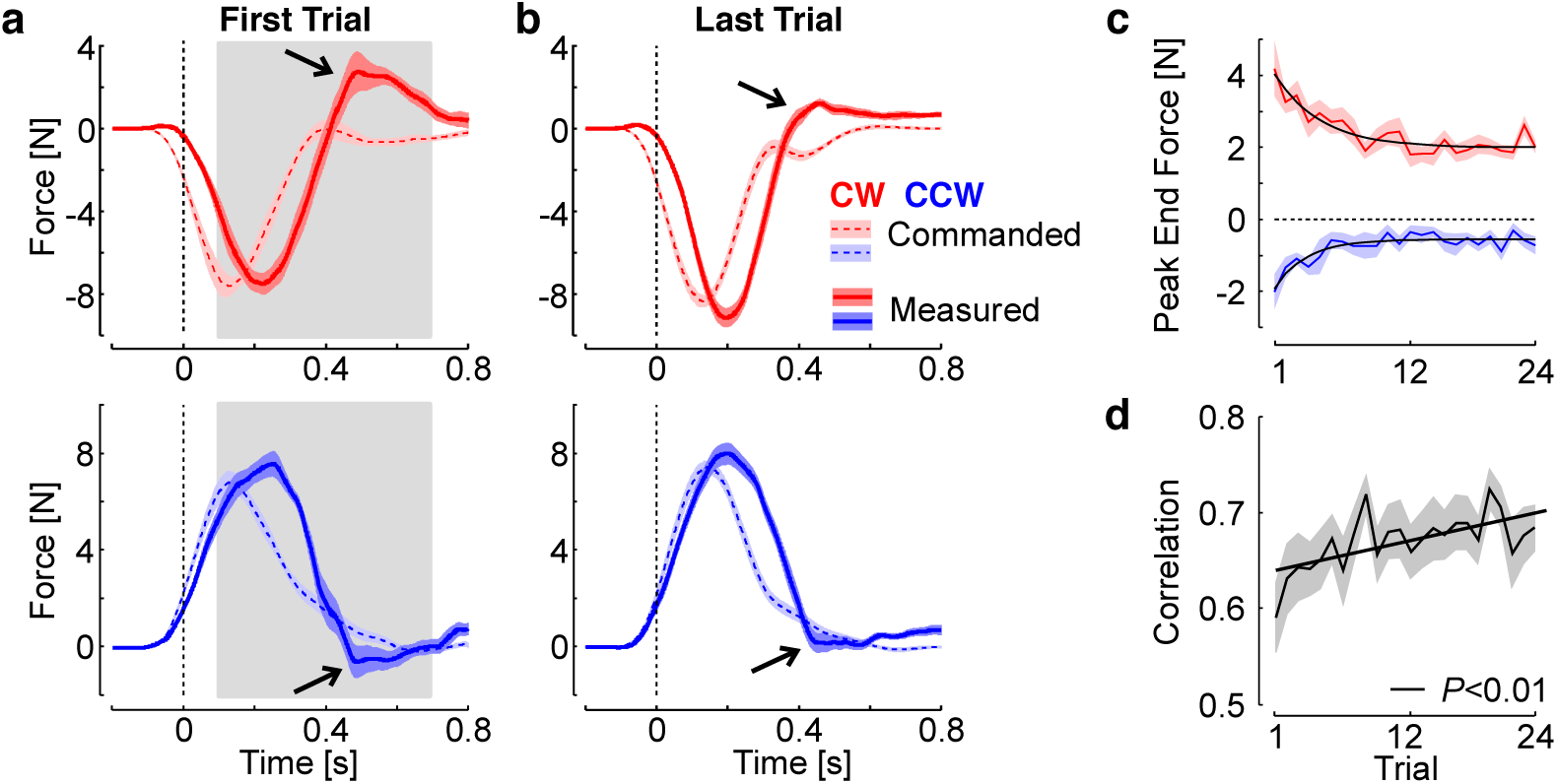
Improvement in Online Corrections. **a:** lateral perturbation force (dashed/light) and measured force (solid/dark) as a function of time for the first force field trial in each direction. **b**: Same as panel a for the last perturbation trial. Shaded areas represent one SEM across participants (n=14) with the same color code as in Fig.1. Note that each force field induced a reaction force with opposite sign, thus the perturbation force was multiplied by −1 for illustration. The reduction in peak end force is highlighted with the black arrows. **c**: Peak end force across force field trials (mean±SEM). The solid traces show the significant exponential decay (*P*<0.005). **d**: Correlation between the commanded force and the measured force during the time interval corresponding to the gray rectangle in panel a (from 100ms to 700ms). The correlations were averaged across participants; the shaded area represents one SEM. The significant linear regression of the correlations across blocks is illustrated.

**Figure 3.**
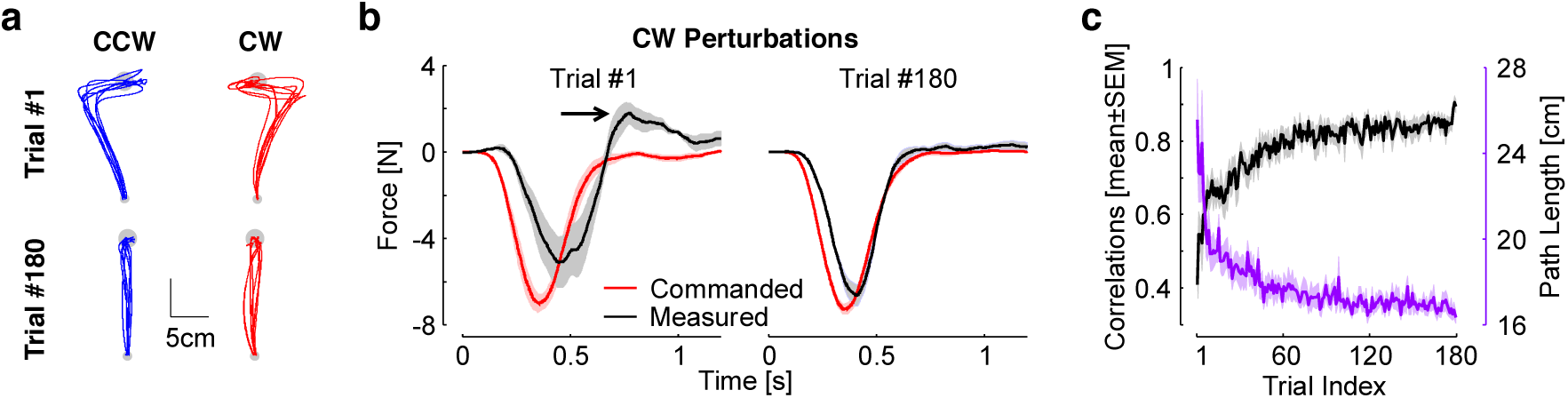
Control experiment. **a.** Hand paths of the first and last trials for each perturbation direction. Displays are individual movements corresponding to one trace for each participant. **b.** Commanded and measured forces for the first and last trials. Shaded areas are one SEM across participants (n=8). **c.** Black: correlations between commanded and measured forces across trials. Purple: Path length across trials (mean±SEM across participants).

The results of Experiment 1 suggested that the controller changed during movement. Indeed, the absence of an effect on the maximum lateral deviation indicated that participants started the movement with a controller that would produce a straight reach path in the absence of any force field (“baseline” controller, *C*(*B*)), and then corrected their movements by using a controller that was partially adapted to the force field (*C*(*F*_*i*_) for the *i*^th^ force field trial). It is important to realize that if the nervous system always switched to the same controller during force field trials, it would have been impossible to identify a change in representation reflecting rapid adaptation, from a standard feedback response. The evidence for rapid adaptation is not based on the feedback response observed during each perturbation trial per se, but on the change in feedback responses across these trials. In other words, the controller (*C*(*F*_*i*_) changed across force field trials (*C*(*F*_*i*_) ≠ (*C*(*F*_*j*_), *i* ≠ *j*). The fact that the perturbations were not anticipated suggests that the adaptive change from (*C*(*B*) to (*C*(*F*_*i*_) during the *i*^th^ force field trial was based on sensory information collected during the ongoing trial. Interestingly the presence of an after-effect indicated that these within-trial changes in control are likely linked to the fast time scales of trial-by-trial learning (Smith et al., 2006).

### Control Experiment

The control experiment was designed to characterize participants’ behavior in a standard trial-by-trial learning and verify that the correlation between the commanded force and the measured force reflected adaptation. The results are shown in Fig. 3. Panel a highlights that the first trials in each force field was strongly perturbed, while the last trials were relatively straight (one trace per participant). Clearly the measured forces presented similarities the ones observed in Fig. 2. For the first trial, the measured force responded late to the force field (black trace lagging the red trace in Fig. 3b, left), which was followed by feedback correction and overcompensation (see black arrow, peak end force). With practice participants learned to anticipate the force field, and the overcompensation disappeared (Fig. 3b, right).

It should be observed that at the end of the 180 trials, the commanded and measured forces are not exactly the same. This could be due to several factors: first the forces linked to the KINARM intrinsic dynamics (*f*_*R*_ in Eq. 2). As well, participants may not have learned to fully compensate for the force field, as the last movements still exhibited some curvature. Nevertheless, the correlations between commanded and measured forces clearly presented an increase across trials typical of a learning curve (Fig. 3c, black). This learning curve paralleled another measure of adaptation based on the path length, which is 15cm for perfectly straight movements (Fig. 3c, purple). We calculated significant exponential fits for both learning curves (*P*<0.005, R^2^= 0.44 for the correlations, and 0.32 for the path length), and observed a slightly faster rate for the path length (CI: [-0.059, −0.043]) in comparison to the correlations (CI: [-0.036, −0.025]), meaning that the path length and correlations approached their asymptotes after 60 and 100 trials, respectively (3 time constants).

These force profiles were broadly similar to those of Experiment 1, with the clear difference that there was no anticipation in the random context of Experiment 1. The decrease in peak terminal force was clearly present in both cases. The correlations at the end are close to 85% on average. The remaining 15% can be ascribed to unmodelled KINARM dynamics (recall that we evaluated an impact of ≤10%, see Methods) and to the fact that adaptation was likely not 100%.

### Experiment 2

The results of Experiment 1 are consistent with adaptive control. However it is possible that participants learned to produce smooth corrective movements by altering movement kinematics without learning about the force field specifically, or by altering the mechanical impedance of the limb without changing their internal model (Burdet et al., 2001). To address this, we sought a more direct link between the online correction and the force field by using a via-point located at two thirds of the reach path (see Methods). We reasoned that if the feedback correction during force field trials reflected an online update of the internal models, then these corrections should evoke almost instantaneous after-effects for the remainder of the movement (from the via point to the goal).

Our results confirmed this prediction. First we reproduced the observations made in the first experiment for the trials without via-point: we found non-significant exponential decay for the maximum lateral deviation, whereas the target overshoot displayed a significant exponential decay across trials (Fig. 4 a-c). The gradual increase in correlation between the commanded force and the applied force was also reproduced in this dataset for the trials without via-point (Fig. 4d). Second, for the via-point trials, we found a systematic after-effect between the via-point and the goal that mirrored the impact of the force field prior to the via-point (Fig. 4e for average hand paths and 4f). We observed that the via-point trials were challenging and many did not reach the goal target on time. We measured a significant increase in success rate across blocks for all types of trials with via-point (rmANOVA, *P* < 0.005 for CW and CCW trials, and *P* = 0.006 for baseline trials), and observed a tendency to reduce overshoot at the via-point similar to the trials without via-point. In the following analyses, we included all trials that stopped at the via-point even if they did not manage to reach the second target.

**Figure 4:**
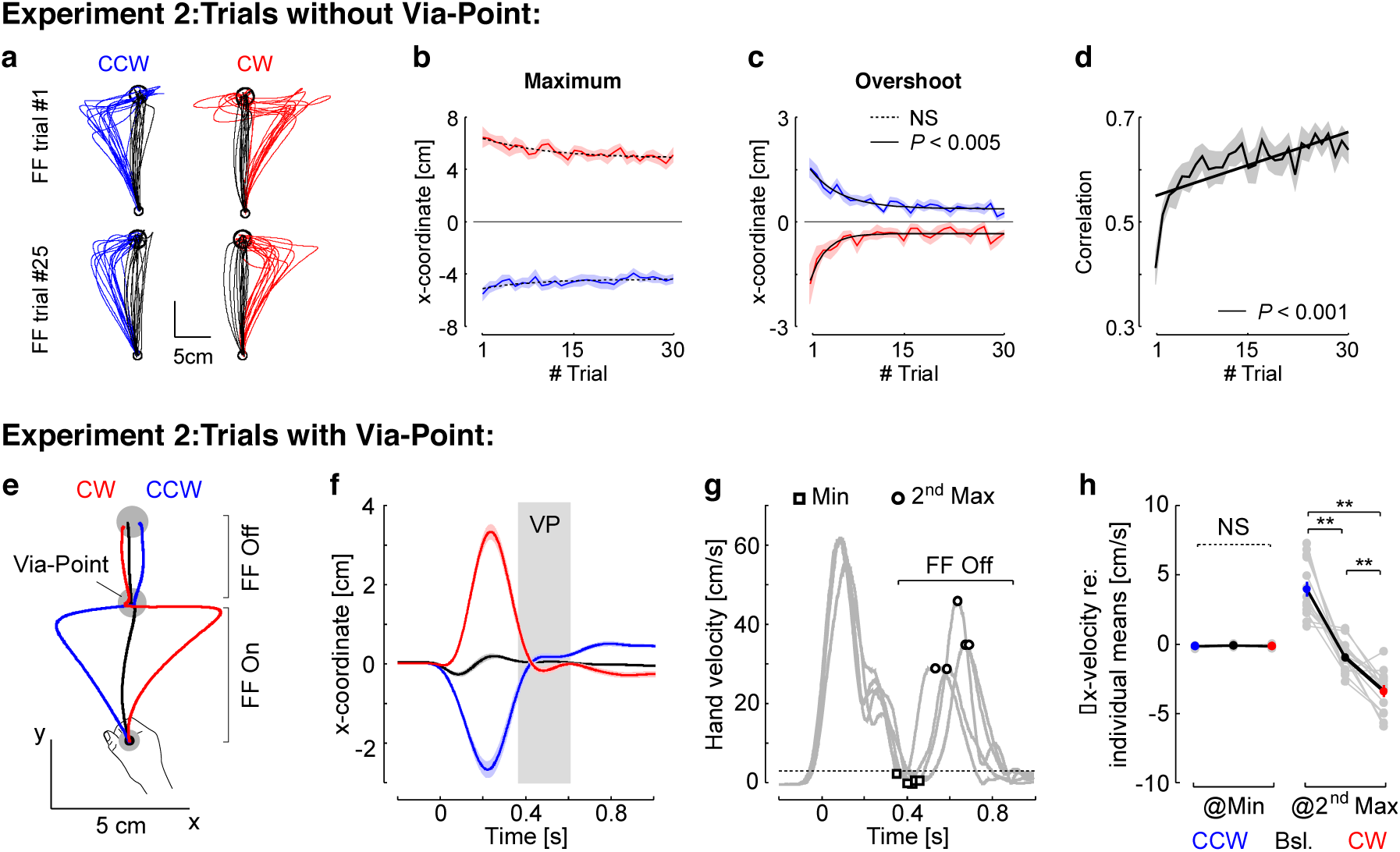
Experiment 2 Behavior. a-c: illustration of the force field trials and behaviour as reported in Fig. 1b-d. d: correlations between the commanded force and the measured force across force field trials similar to Fig. 2d. e: Average hand traces from Experiment 2 for trials with a via-point with the same color code as in Experiment 1 (black: null-field, red: CW, blue: CCW). For all trials, the second portion of the reach, from the via-point to the goal, was performed without force field (FF Off). f: Average x-coordinate as a function of time across participants (n=14). Shaded areas represent one SEM across participants. The gray rectangle illustrates the approximate amount of time spent within the via-point. Observe the displacement following the via-point in the opposite direction as the perturbation-related movement. g: Illustration of the trial-by-trial extraction of parameters on five randomly selected trials from one participant. The force field was turned off when the hand speed dropped below 3 cm/s (dashed trace). The parameters extracted were the lateral hand velocity at the moment of the hand speed minimum (squares), and the lateral hand velocity at the moment of the second hand speed maximum (open discs). h: Lateral hand velocity at the moment of hand speed minimum or at the moment of the second hand speed maximum averaged across trials for each participant (gray dots), and averaged across participants (blue, black and red, mean±SEM). Individual means across the three trial types were subtracted from the data for illustration purpose. The stars represent significant pair-wise differences at the level *P*<0.005 with Bonferroni correction for multiple comparisons.

Stopping at the via-point was necessary for two reasons: first it allowed us to turn off the force field unnoticeably because the speed was close to zero. Second, it also ensured that the after-effects were not due to momentum following the online correction prior to the via-point, because the speed was very close to zero at the via-point. To further verify that it was not an effect of momentum, we extracted the minimum lateral hand velocity in the via-point and the lateral hand velocity at the second peak (Fig. 4g: squares and open discs on exemplar traces). As expected, there was no difference across clockwise, baseline, and counterclockwise trials for the minimum velocity (Fig. 4h, rmANOVA, *F*_(2,39)_=1.13, *P*=0.33), which confirmed that there was no difference in momentum across trials at the via-point. The same analysis with the norm of hand velocity in place of lateral velocity gave identical results. In contrast, the lateral hand velocity measured at the second peak hand speed displayed a very strong modulation consistent with an after-effect (Fig. 4h, *F*_(2,39)_ = 50.59, *P*<10^−5^). In addition, we observed that 12 out of the 14 participants showed the same modulation when comparing the distributions of individual trials across CW and CCW perturbations (Wilcoxon ranksum test, *P*<0.05). Thus, the after-effects between the via-point and the goal were due to a re-acceleration in the direction opposite to the force field experienced during the first part of the trial.

By forcing participants to stop at the via-point, we controlled experimentally for the effect of momentum prior to the via-point. We conducted additional analyses to address the possible influence of hand kinematics prior to the via-point based on statistical modeling. First, we regressed the lateral velocity at the second peak as a function of lateral velocity extracted at the first velocity peak to address the influence of the perturbation experienced prior to the via point (Fig. 5a). We fitted a linear mixed model on individual trials from all participants while including a random intercept to account for idiosyncrasy. The regression was computed first on the baseline trials only, and we found no significant correlation between the lateral velocity before and after the via-point (Fig. 5b, dashed line, *F*_(344)_ = 0.31, *P* = 0.75). Second we fitted another mixed model including force field trials and found a significant relationship between velocity prior and after the via-point (Fig. 5b, gray line, *F*_(888)_ = 16.09, *P*< 0.0001). We then fitted a third model that included an interaction between the first hand velocity, and the categories of trials (CCW, CW, baseline). The analysis of covariance with this model revealed again a strong influence of the first velocity peak (*F*_(1,886)_=267, *P*<0.0001) as well as a significant interaction between the first velocity peak and the categories (*F*_(2,886)_16, *P*<0.0001). The comparison of the two models based on BIC indicated that the best model was the one including the categories as a factor. Thus, the modulation of hand velocity after the via-point was best accounted for by a statistical model that included the type of perturbations, indicating that the modulation of hand velocity was not simply accounted for by movement kinematics prior to the via-point.

**Figure 5.**
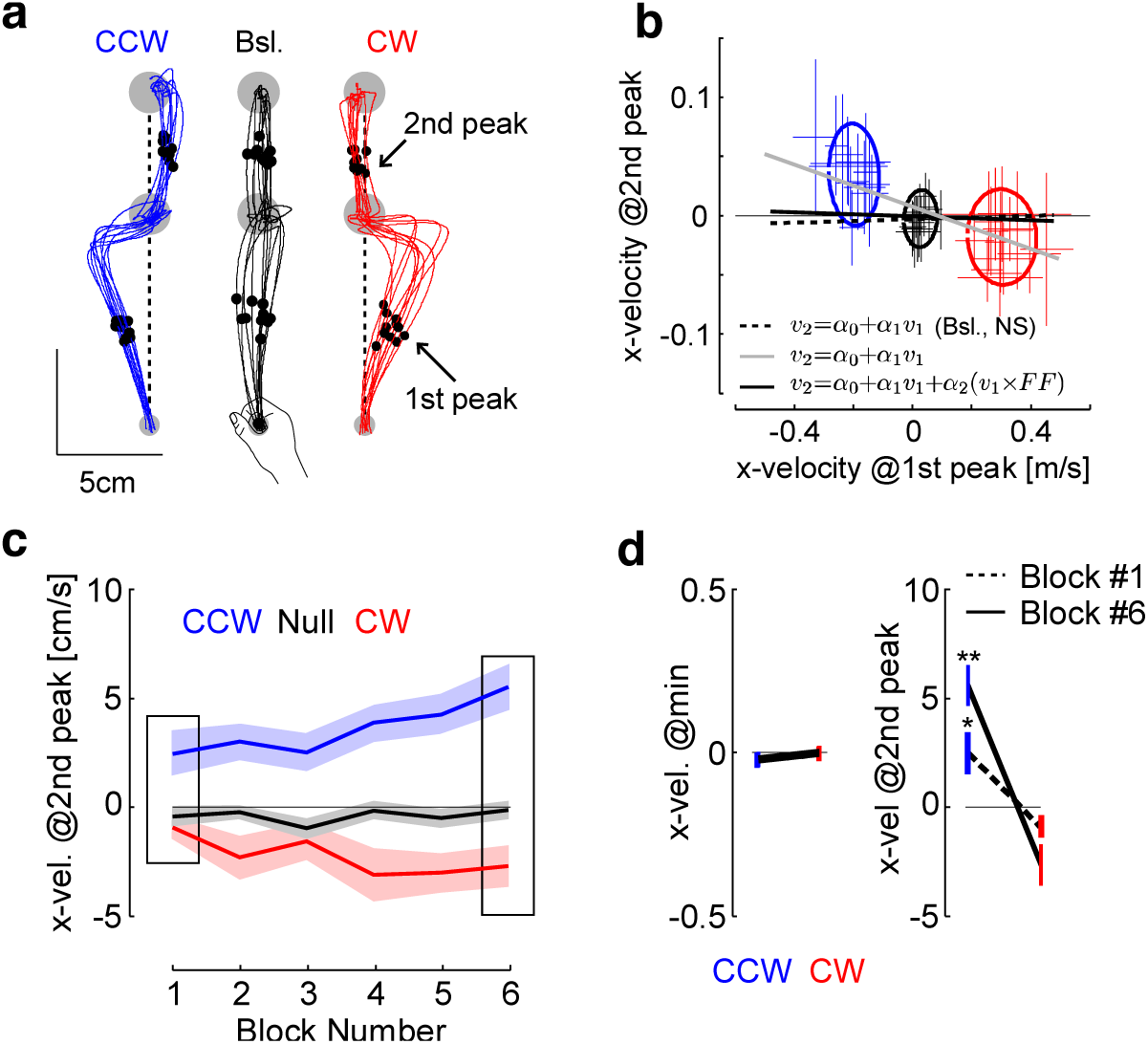
Kinematics after the via-point. a. Extracted parameters. The lateral velocity (x-direction) was extracted at the moment of the first and second peaks of hand speed prior and after the via-point, respectively (dots and arrows). Vertical dashed lines were added to highlight the systematic deviation opposite to the perturbation during force field trials. We selected 10 traces randomly per condition from one representative participant. b. Lateral velocity at the second peak as a function of the lateral velocity at the first peak. Crosses are mean ± standard deviation across trials for each participant (n=14). Dispersion ellipses illustrate one standard deviation along the main variance axes (all trials pooled together). The three statistical models are illustrated with thick lines: baseline trials only (not significant), all trials (gray), and all trials with categories as factor (black plus average of the categories represented with the ellipses). The thin black line is the value of 0 displayed for illustration. Participants were included as random factor. c: Lateral hand velocity measured at the second peak hand speed following CCW and CW force-fields (red and blue, respectively), and baseline trials (black) across blocks. Data are from 11 participants who completed successful trials in each block. The insets highlight the first and last block. d: Group data mean±SEM in the 1^st^ and 6^th^ block for the lateral hand velocities at the minimum and at the second peak (n=11 in both cases). Observe the two different scales. One (two) star(s) highlight significant difference based on paired t-test at the level *P*<0.05 (*P* < 0.005).

Second, we extracted the evolution of trials across the blocks and observed a significant interaction between the force field and the block number across blocks #1 and #6 (Fig. 5c and d, *F*_(2,20)_ = 4.08, *P*=0.032). This analysis was restricted to the data from 11 participants who completed at least one successful via-point trial in these blocks to balance the statistical test. Strikingly, the modulation of hand velocity following the via-point was already present in the first block (Fig. 5d, paired t-test on data from block 1: t_10_ = 2.7, *P* = 0.0106, effect size: 0.98, data from block 6: t_10_ = 6.3, *P*<0.0001, effect size: 1.56). Again we verified that there was no effect of the force-field and no interaction between force field and block number for minimum hand speed within the via-point (*F*_(2,20)_<1.7, *P*>0.2). Third, we assessed whether the lateral hand velocity prior to the via-point exhibited changes across the blocks and found no significant change for both CW and CCW perturbations (*F*_(5,72)_<1.69, *P*>0.1). In all, these analyses indicated that there was no kinematic effect from the first part of the movement, or residual momentum at the via-point, that could be statistically linked to the lateral hand velocity after the via-point, and to its evolution across blocks.

The presence of a significant modulation in the first block indicated that online adaptation occurred during the first few trials (there were 5 trials with via-point and switching-off of the force field per direction and per block). This striking result warranted further investigation: can these adjustments occur within the very first trial? Our dataset cannot provide a definitive answer because, due to randomization, some via-point trials were preceded by force field trials without via-point for some participants. Nevertheless, we used a statistical model of the modulation of lateral velocity as a function of trial number to obtain the beginning of an answer.

We first changed the sign of velocity during CW trials such that an after effect corresponded to a positive velocity for all trials. Then, we accounted for idiosyncrasy by including participants as a random factor in a mixed linear model. We found a significant positive correlation between the lateral hand velocity and the trial index. Most importantly, the regression included an offset that was significantly distinct from 0. The intercept value was 0.02ms^-1^, and the *P-*values for the slope and intercept were 0.0026 and less than 10^−4^, respectively. The same regression performed on the baseline trials, where no force field was applied prior to the via-point, revealed an offset of −0.004, which was not significant (*P* = 0.15). Thus, we can reject the hypothesis that the lateral hand velocity was zero in the very first trial. In other words, our dataset supports the hypothesis that online learning may have occurred during the very first trial.

To summarize, Experiment 2 showed that hand trajectories after the via-point were compatible with an after effect evoked by the presence of a force field prior to the via-point. The experimental design and statistical models were used to control for momentum and for kinematic effects. Furthermore, statistical modeling also indicated that the reported adjustments occurred within the first few force field trials. A priori, it is possible that participants used three internal models, one for null-field trials and one per force-field direction, and switched between them dependent on the ongoing movement. However, this explanation does not easily account for the fact that participants never experienced force field disturbances prior to their participation, and thus never adapted to these perturbations. Such a switch within a movement is consistent with adaptive control, but it is not clear how they could switch between models that they did not previously acquire, in particular during the first few force field trials exhibiting improvements in online corrections. A more compelling explanation in our opinion, which explains the results of the two experiments as well as the increase in modulation reported in Experiment 2, is the hypothesis that the nervous system tracks model parameters online. This is illustrated in the next section.

### Adaptive Control Model

Our behavioral results highlighted parallels with motor adaptation in spite of the fact that perturbations were not anticipated: the reduction in peak end force, the increase in correlation, and the after effects following the via-points. We show below that these observations can be explained in the framework of adaptive control. The key point is to explain how participants were able to adapt their online response to unexpected force field without anticipation. The controller is composed of a feedback control loop that corrects for perturbations to the state of the limb (Fig. 6a: state-feedback, solid). This loop involves a state estimator through an observer (e.g. Kalman filter for linear systems), and a controller that generates motor commands.

**Figure 6.**
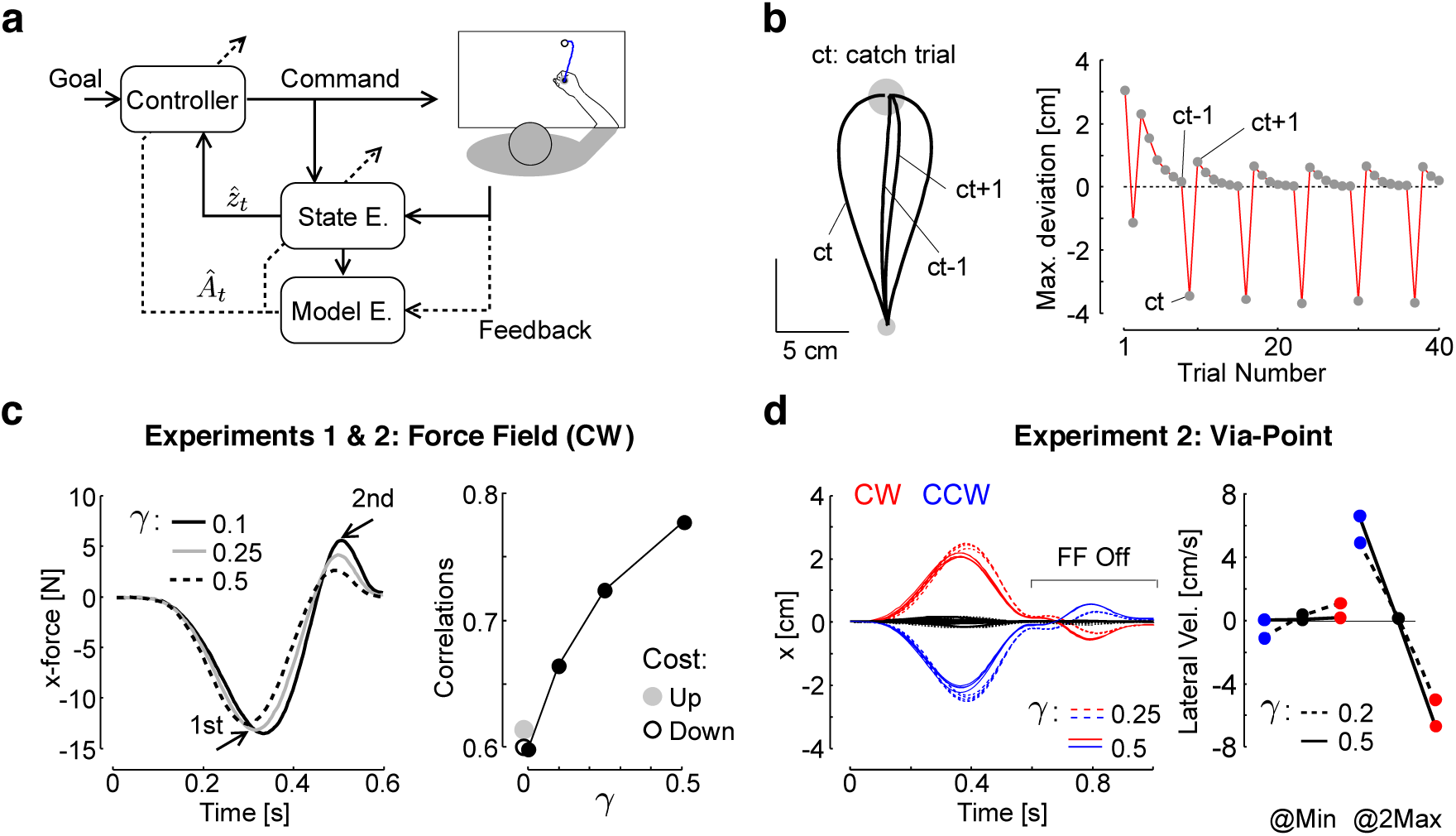
Model Simulations. a: Schematic illustration of an adaptive controller: sensory feedback is used to estimate both the state of the system and the model, which in turn updates the controller and estimator online. The solid arrows represent the parametric state-feedback control loop, and the dashed arrows represent the real-time learning and update of model parameters. b: Reproduction of a standard learning experiment (hand paths and maximum lateral deviation across trials), with mirror-image catch trials (ct), and unlearning following the catch trials (difference between ct-1 and ct+1). The trial-by-trial changes were reproduced with online computations exclusively (inspired by Thoroughman and Shadmehr (Thoroughman and Shadmehr, 2000)). c: Left: Simulation of the results from Experiment 1 with three distinct values of online learning rate (*γ*: 0.1 in black, 0.25 in gray, and 0.5 in dashed), which reduces the second peak hand force while the first displays smaller changes across the tested range of *γ*. The change in peak force relative to the mean across simulations was less than 2N for the first peak, while the second peak displayed changes ∼4N. Right: Correlations between simulated perturbation force and the force produced by the controller. Black dots are the results of the adaptive control model with tested values of *γ* (0, 0.1, 0.25, and 0.5). Gray and open dots are the results of the model with time-evolving cost-function (both increase and decrease). d: Simulations of behavior from Experiment 2, the force field was turned off at the via-point (t = 0.6s), and the second portion of the reach displays an after effect (see Methods). The inset highlights the lateral velocity measured at the minimum and at the second peak hand speed as for experimental data. The traces were simulated for distinct values of online learning rate (0.25 in dashed, and 0.5 in solid).

The second component is the adaptive loop, which performs learning (Fig. 6a: outer loop, dashed). The key idea is that the state feedback control loop is parameterized in a way that accounts for limb and environmental dynamics (Crevecoeur and Kurtzer, 2018), and the function of the adaptive loop is to adjust the parameterization of the state-feedback control loop based on current sensory data and motor commands (Bitmead et al., 1990; Ioannou and J, 1996; Fortney and Tweed, 2012). Classical studies have assumed that the parameterization of the state feedback control loop is modified on a trial-by-trial basis (Shadmehr and Mussa-Ivaldi, 1994; Thoroughman and Shadmehr, 2000; Milner and Franklin, 2005; Smith et al., 2006). In addition, previous modeling work highlighted that there are multiple timescales in motor adaptation (Smith et al., 2006; Kording et al., 2007). We suggest complementing this theory by considering that the fast time scale can be in fact shorter than a trial time. This is mathematically possible: in the framework of adaptive control, learning occurs at each time step. We show that behaviorally, this model accommodates our observations as well as trial-by-trial learning assuming that the acquired representation within a movement carries over to the next one.

Similar to trial-by-trial adaptation models, parameter tracking is indirect because there is no measurement of the unknown parameters, thus motor commands and sensory feedback must be used together to deduce the underlying dynamics. The simulations performed in this study were based on iterative least square identification (LS) (Bitmead et al., 1990), but other techniques such as expectation-maximization may be considered (Ghahramani and Hinton, 1996). However, it is important to note that the increase in after-effect observed across the blocks of Experiment 2 required a degree of freedom in the model to account for trial-by-trial modulation, which is why we privileged the LS formalism as it includes such a parameter by design (online learning rate, *γ*). The implication for our interpretation and its limitations are addressed in the Discussion section.

Using this framework, we reproduced standard learning curves in computer simulations, with exponential decay of the lateral hand displacement, mirror-image catch trials or after effects as documented in Experiment 1 (Fig. 1 e-g), and unlearning following the catch trials as previously reported in human learning experiments (Fig. 6b) (Thoroughman and Shadmehr, 2000). These simulations assumed that the partially corrected model within a movement was used in the next one. This model also reproduced the reduction in peak end force occurring near the end of movement as observed in Experiment 1 by assuming that the online learning rate increased across force field trials (Fig. 6c, left), that is the online adjustments were smaller in the first trials than in the last trials. This idea is consistent with the presence of savings, characterizing a faster re-learning upon exposure to a previously experienced environment (Smith et al., 2006; Gonzalez Castro et al., 2014; Shadmehr and Brashers-Krug, 1997; Caithness et al., 2004; Overduin et al., 2006; Coltman et al., 2019; Nguyen et al., 2019). This assumption was captured here by an increase in online learning rate across simulations (*γ* parameter, note that *γ* was fixed within each trial).

We compared the simulated behavior of Experiment 1 to an alternative hypothesis in which there is no online changes in the representation of the reach dynamics (learning rate *γ* = 0), and in which control gains are re-adjusted at each time step by increasing or decreasing the cost parameters within movements. First an increase in cost was clearly incompatible with the data since it generated an increase in peak end force, whereas we measured a strong exponential decay across force field trials. In contrast, the decrease in cost produced a reduction in peak terminal force, but also in the first peak later force (in absolute value), which was again not consistent with our dataset. In addition, the model with reduction in cost-function produced larger lateral displacements due to a reduced penalty on state deviation, which we did not observe empirically (not shown).

The strongest argument in support to the adaptive control model was obtained by calculating the correlations between simulated forces and the controller force. The measured force in the data is the force applied by the hand to the handle. It can be equated to the force applied by the controller to the point mass. In theory it only depends on the control vector, whereas in practice it also depends on the arm passive dynamics. Thus the comparison has limitations but it is valid for the purpose of this analysis. Recall that we neglected the robot dynamics on the basis that it had a small impact on the estimation of hand acceleration. We found that by increasing the value of the online learning rate, the correlations increased in a range compatible with the data (Fig. 6c, black dots, compare with Figs 2d and 4d). In contrast, the models with time evolving cost-functions did not produce any change in correlation (Fig. 6c, gray and open dots), which suggested that the change in correlation was a signature of adapting the internal model.

Lastly, it is important to highlight that models based on changes in impedance control or time-evolving cost functions without any change in online representation would not produce any after-effect after force field trials (Fig. 1, f and g) or after the via-point (Fig. 4h), since these would assume no change in expected dynamics. In contrast, the adaptive control model also reproduced the after effect observed following a single force field trial (Fig. 1g and 6b), as well as the modulation of lateral hand velocities following the via-point (Fig. 6d). The adjustment of online learning rate based on the assumption of savings was also necessary in these simulations to account for the increase in lateral hand velocity across the blocks (Figs. 5c and 6d). In all, adaptive control combined with the possibility of an increase in online learning rate across trials explains the modulation of hand force and adaptation of feedback responses during movement (Figs. 2, 3, 4d, and 5c), rapid adaptation following via-point trials (Figs. 4e-h and 6d), and trial-by-trial learning.

## Discussion

Controlling movements in the presence of model errors is a challenging task for robotics as well as for the nervous system. However healthy humans can handle model errors like those arising when a force field is introduced experimentally during reaching. To understand this, we explored the possibility that the nervous system reduces the impact of model errors within a movement following the principles of adaptive control. We found that participants’ feedback corrections to unexpected force fields gradually improved (Experiments 1 and 2), and evoked after-effects in an ongoing sequence of movements (Experiment 2) within 500ms or less. We showed that these effects, as well as single rate exponential learning curves, were captured in an adaptive control model in which motor commands, state estimates, and sensory feedback are used to update the internal models of dynamics online. We suggest that this model is a powerful candidate for linking movement execution with the fast time scales of trial-by-trial motor learning.

First this model is built on current theories of sensorimotor coordination (Todorov and Jordan, 2002). There is no disagreement between adaptive control and stochastic optimal control (LQG, (Todorov, 2005)), but instead there is complementarity between these models. Stochastic optimal control is based on the assumption that the system dynamics are known exactly, and that the process disturbance follows a known Gaussian distribution (Astrom, 1970). The unexpected introduction of a force field introduces a fixed error in the model parameters, which is not a stochastic disturbance. In theory, such disturbances warrant the use of a control design that explicitly considers the presence of errors in the model parameters. Adaptive control aims at solving this problem by learning about the dynamics during movement.

Real-time adaptive control also stands in agreement with learning and acquisition of motor skills over longer timescales. Our analyses suggested in Fig. 6b were based on the idea that if the change in representation acquired within a movement carried over to the next movement, then we can reproduce exponential decay, after effects and unlearning following catch trials. This possibility is not in conflict with the fact that there exist slower changes in the nervous system supporting consolidation and long-term retention. In fact, adaptive control simply consists in adding a timescale shorter than the trial time to current models (Smith et al., 2006; Kording et al., 2007), with the major implication that, by being faster than a trial, adaptation becomes available to complement feedback control. The theory of adaptive control was put forward to highlight that this was mathematically possible, and theories are there to be challenged. In our opinion, this theory is the simplest combination of control and adaptation models: it adds adaptation to current models of movement execution by considering a fast time scale in a standard learning model.

Of course, it is important to consider candidate alternative mechanisms. The main findings that require revision of the theory are that participants made rapid, field-specific adjustments to their motor output in an unpredictable task environment. The main pieces of evidence that point to rapid adaptation are (1) the increase in correlation between commanded and applied forces also observed in trial-by-trial adaptation (Figs. 2d, 3c, and 4d); and (2) the presence of an after effect within 500ms (Fig. 4f). The value of the adaptive control model is to provide a mathematical framework in which these observations are expected, and which combines current models of learning and control with the single addition of a timescale faster than a trial.

The shortcomings in our interpretations are the following. First, our proposed model required a change in online learning rate (*γ*) to accommodate the improvement in online feedback control in the absence of anticipation and without previously acquired internal models. This assumption was based in part on the observation that the learning rate can change dependent on the context (Gonzalez Castro et al., 2014), and on the hypothesis of savings characterized by the ability to re-learn faster. This hypothesis has been documented in many tasks including adaptation to force fields (Shadmehr and Brashers-Krug, 1997; Caithness et al., 2004; Overduin et al., 2006; Coltman et al., 2019; Nguyen et al., 2019). Here we used the same concept applied to the online learning rate. This is of course a strong assumption that requires further validation. Second, we drew a parallel between the fact that feedback responses improved across force field trials in Experiments 1 and 2, and that the magnitude of the after effect after the via-point increased. These two observations are consistent with the increase in online learning rate. However, they also exhibited different rates, indicating that several mechanisms could be at play.

Alternative models must be also considered. A first is the possibility that participants used co-contraction to modulate the mechanical impedance of their limb and counter unpredictable or unstable loads (Hogan, 1984; Burdet et al., 2001; Milner and Franklin, 2005). We do not reject this possibility, but must underline that it does not explain all the data. In particular, the results from Experiment 2 emphasized an after effect after the via-point, which is not expected if participants only made their arm mechanically rigid. In such case one expects a straighter hand path instead of a deviation opposite to the force field. A second argument that does not favor impedance control was addressed in the model: by changing the cost-function during movement, we effectively increased the elastic and damping components of control gains, as there was no delay in the model. We showed in the (simplest) settings of our model that such a strategy would not produce an increase in correlation compatible with the one observed in the data (Fig. 6c). A third objection is based on a recent observation made in a similar unpredictable environment (Crevecoeur et al., 2019). In this case, participants co-contracted spontaneously following force field trials, which reduced lateral deviations for subsequent perturbations. This has a small but measurable impact (∼10%), which likely explained the small reduction observed in Figs. 1c and 4b, but the questions stand out as how participants then managed to fall on target while reducing the force applied to the handle. Thus, as first approximation this model does not stand as the strongest candidate, but a more detailed account of the possible role of impedance control in such task is warranted.

Another possibility is that participants switched between controllers corresponding to CW or CCW force fields shortly following movement onset. In fact, such a switching between controllers is a form of adaptive control with a limited, or finite set of representations. In this framework, contextual cues such as feedback error signals could be used to switch between multiple internal models as suggested in the MOSAIC framework (Haruno et al., 2001). Both switching between internal models and adaptive control include a change in representation within movement, which is compatible with our data. The open question is whether the change was discrete (between two or more internal models), or continuous (as in online parameter tracking). Two possible objections are that the internal models corresponding CW and CCW perturbations were not previously acquired, and that the change in control seemed more gradual than discrete, as it mostly impacted control near the end of the movement.

In fact, although possible, this interpretation would disagree with a large body of literature showing that exposition to opposite velocity-dependent force fields within short times leads to partial to total interference (Brashers-Krug et al., 1996; Gandolfo et al., 1996; Hwang et al., 2003). Based on these results we would thus expect that participants could not have acquired an internal model for each force field. It was then shown that distinct planning conditions were critical to learn opposite perturbations, such as explicit cues (Wada et al., 2003; Osu et al., 2004; Addou et al., 2011), distinct representations of movements (Hirashima and Nozaki, 2012), or even distinct prior or follow-through movements associated with the same movement (Howard et al., 2012; Sheahan et al., 2016). In our study, there was no cue given about the upcoming perturbation and planning was likely identical across baseline and force field trials. Thus, it was the physical state of the limb during movement that evoked the changes in control during movements. Part of the discrepancy may simply be that adaptation became visible near the end of movements, where classical studies would focus on metric such as maximum lateral deviation, based on which we would also conclude that there was no learning (e.g. Fig. 1c). But interference was not confined to the metric chosen; it also impacted learning curves over several trials. Adaptive control may solve this apparent contradiction: interference does occur in our model as perturbations are applied randomly (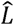 on average is 0), but the online parameter tracking enabled adapted corrections.

The question arises as whether the after effect after the via-point could be considered as a second sub-movement. In this case the after effect documented after the via-point would not be distinct from standard after effects. This was the point of our developments: we wanted to highlight standard after effects. Our key contribution was to show that the second sub-movement (from the via-point to the target) was re-planned, or changed, within 500ms (Fig. 4), since the same movement without force field was on average straight. This rapid change was fast enough to influence control for standard force field trials, and faster than previously identified time scales of motor adaptation.

It remains unclear how quickly these adjustments occurred within movements, and this represents an important but challenging question for future research. Anatomically, long-latency corrections for perturbation to the limb (∼60ms) are supported by a distributed network through cerebellum (Hore and Flament, 1988), primary somatosensory area, motor and pre-motor cortices, and parietal cortex (Pruszynski et al., 2011; Omrani et al., 2016; Scott, 2016). This network clearly overlaps with the main regions associated with short-term plasticity (Dayan and Cohen, 2011). Behaviorally, long-latency feedback exhibits exquisite sensitivity to even very small disturbances (Crevecoeur et al., 2012). Thus, the network of brain regions engaged in trial-to-trial learning is also likely recruited very early during force field trials, and it is conceivable that rapid adjustments occurred within this rapid pathway.

Computationally, however, an online change in internal models involves an indirect mapping of sensory feedback into model updates, in addition to corresponding changes in the neural controller. This computational load may require longer processing times associated with model updates in comparison with feedback responses following learned perturbations to the limb or to the movement goal. As an absolute upper bound, we may consider that adjustments in Experiment 2 were performed between reach onset and the moment when participants exited the via-point, which was around 500ms. Similarly, visual inspection of the forces from Experiment 1 indicates that changes could have occurred earlier. However, the precise timing of change in feedback correction cannot be unambiguously determined from the resultant force since this variable involves both intrinsic limb properties and neural feedback. We expect that future work measure the latency of this outer loop more accurately based on muscle recordings.

Another clear challenge is to unravel the underlying neural mechanism. One key question is to understand how neural circuits generated a near instantaneous after effect like those observed in Experiment 2. Indeed, at the via-point with near-zero velocity across all trials, there was no motor error to correct; yet the subsequent motor output mirrored the previously encountered disturbances. One possibility to account for this result is through synaptic plasticity (Dayan and Cohen, 2011): that is the gain of the same sensorimotor loop changes following re-afference of sensory prediction errors, and associated changes in synaptic strength. However, the fast time scales of synaptic plasticity documented in cerebellar-dependent adaptation remain longer than a trial time (Raymond and Medina, 2018). Another possibility is that the unexpected force field generated different neural trajectories across force field trials, possibly exploiting dimensions that do not directly influence the motor output (Kaufman et al., 2014; Stavisky et al., 2017). Thus, a similar perturbation can produce distinct motor outputs by tuning feedback circuits. In theory such mechanism does not necessarily engage changes in synaptic weights, which is computationally advantageous (Fortney and Tweed, 2012). Furthermore, changes in the null dimension of the neural manifolds were recently reported as neural-basis of trial-by trial learning (Perich et al., 2018), which makes such a mechanism a very strong candidate to support within-trial adaptation of movement representations. Unraveling the latency of the model updates will likely set physiological constraints on the candidate neural underpinnings.

Finally, our study also opens questions from a theoretical standpoint. Indeed, we must recall that our model was excessively simple: we considered the translation of a point-mass in the plane with full state information (see Methods). We thus expect that future modeling work unveil the computational challenges that arise when considering more realistic models of the neuro-musculoskeletal system, including nonlinear dynamics, several unknown parameters, sensorimotor noise, and temporal delays.

## References

Addou T, Krouchev N, Kalaska JF (2011) Colored context cues can facilitate the ability to learn and to switch between multiple dynamical force fields. J Neurophysiol 106:163–183.

Astrom KJ (1970) Introduction to stochastic control theory. New York: Academic Press.

Bitmead RR, Gevers M, Wertz V (1990) Adaptive optimal control : the thinking man’s GPC. New York: Prentice Hall.

Brashers-Krug T, Shadmehr R, Bizzi E (1996) Consolidation in human motor memory. Nature 382:252–255.

Braun DA, Aertsen A, Wolpert DM, Mehring C (2009) Learning Optimal Adaptation Strategies in Unpredictable Motor Tasks. Journal of Neuroscience 29:6472–6478.

Burdet E, Osu R, Franklin DW, Milner TE, Kawato M (2001) The central nervous system stabilizes unstable dynamics by learning optimal impedance. Nature 414:446–449.

Burdet E, Osu R, Franklin DW, Yoshioka T, Milner TE, Kawato M (2000) A method for measuring endpoint stiffness during multi-joint arm movements. Journal of Biomechanics 33:1705–1709.

Caithness G, Osu R, Bays P, Chase H, Klassen J, Kawato M, Wolpert DM, Flanagan JR (2004) Failure to consolidate the consolidation theory of learning for sensorimotor adaptation tasks. Journal of Neuroscience 24:8662–8671.

Cluff Y, Scott SH (2013) Rapid feedback responses correlate with reach adaptation and properties of novel upper limb loads. Journal of Neuroscience 33:15903–15914.

Coltman SK, Cashaback JGA, Gribble PL (2019) Both fast and slow learning processes contribute to savings following sensorimotor adaptation. J Neurophysiol 121:1575–1583.

Crevecoeur F, Scott SH (2013) Priors Engaged in Long-Latency Responses to Mechanical Perturbations Suggest a Rapid Update in State Estimation. Plos Computational Biology 9.

Crevecoeur F, Scott SH (2014) Beyond Muscles Stiffness: Importance of State Estimation to Account for Very Fast Motor Corrections. PLoS Computational Biology 10:e1003869.

Crevecoeur F, Kurtzer IL (2018) Long-latency reflexes for inter-effector coordination reflect a continuous state-feedback controller. J Neurophysiol.

Crevecoeur F, Kurtzer I, Scott SH (2012) Fast corrective responses are evoked by perturbations approaching the natural variability of posture and movement tasks. J Neurophysiol 107:2821–2832.

Crevecoeur F, Scott SH, Cluff T (2019) Robust control in human reaching movements: a model-free strategy to compesate for unpredictable disturbances. Journal of Neuroscience.

Dayan E, Cohen LG (2011) Neuroplasticity subserving motor skill learning. Neuron 72:443–454.

Diedrichsen J (2007) Optimal task-dependent changes of bimanual feedback control and adaptation. Curr Biol 17:1675–1679.

Diedrichsen J, White O, Newman D, Lally N (2010) Use-dependent and error-based learning of motor behaviors. J Neurosci 30:5159–5166.

Fortney K, Tweed DB (2012) Computational Advantages of Reverberating Loops for Sensorimotor Learning. Neural Computation 24:611–634.

Franklin DW, Wolpert DM (2011) Computational Mechanisms of Sensorimotor Control. Neuron 72:425–442.

Gandolfo F, Mussa-Ivaldi FA, Bizzi E (1996) Motor learning by field approximation. Proc Natl Acad Sci U S A 93:3843–3846.

Ghahramani Z, Hinton GE (1996) Parameter estimation for linear dynamical systems. Technical Report CRG-TR-96-2, University of Toronto, Dept of Computer Sceince.

Gonzalez Castro LN, Hadjiosif AM, Hemphill MA, Smith MA (2014) Environmental consistency determines the rate of motor adaptation. Curr Biol 24:1050–1061.

Haruno M, Wolpert DM, Kawato M (2001) Mosaic model for sensorimotor learning and control. Neural Comput 13:2201–2220.

Hirashima M, Nozaki D (2012) Distinct Motor Plans Form and Retrieve Distinct Motor Memories for Physically Identical Movements. Current Biology 22:432–436.

Hogan N (1984) Adatpive-control of mechanical impdance by coactivation of antagonist muscles. Ieee Transactions on Automatic Control 29:681–690.

Hore J, Flament D (1988) Changes in motor cortex neural discharge associated with the development of cerebellar limb ataxia. J Neurophysiol 60:1285–1302.

Howard IS, Ingram JN, Franklin DW, Wolpert DM (2012) Gone in 0.6 seconds: the encoding of motor memories depends on recent sensorimotor states. J Neurosci 32:12756–12768.

Hwang EJ, Donchin O, Smith MA, Shadmehr R (2003) A gain-field encoding of limb position and velocity in the internal model of arm dynamics. PLoS Biol 1:E25.

Ioannou P, J S (1996) Robust Adaptive Control: Prentice Hall Inc.

Izawa J, Rane T, Donchin O, Shadmehr R (2008) Motor adaptation as a process of reoptimization. Journal of Neuroscience 28:2883–2891.

Kaufman MT, Churchland MM, Ryu SI, Shenoy KV (2014) Cortical activity in the null space: permitting preparation without movement. Nat Neurosci 17:440–448.

Kording KP, Tenenbaum JB, Shadmehr R (2007) The dynamics of memory as a consequence of optimal adaptation to a changing body. Nat Neurosci 10:779–786.

Krakauer JW, Shadmehr R (2006) Consolidation of motor memory. Trends Neurosci 29:58–64.

Krakauer JW, Ghilardi MF, Ghez C (1999) Independent learning of internal models for kinematic and dynamic control of reaching. Nature Neuroscience 2:1026–1031.

Kurtzer IL, Pruszynski JA, Scott SH (2008) Long-latency reflexes of the human arm reflect an internal model of limb dynamics. Current Biology 18:449–453.

Lackner JR, DiZio P (1994) Rapid adaptation to Coriolis-force perturbations of arm trajectory. J Neurophysiol 72:299–313.

Lackner JR, DiZio P (2005) Motor control and learning in altered dynamic environments. Current Opinion in Neurobiology 15:653–659.

Laird NM, Ware JH (1982) Random-effects models for longitudinal data. Biometrics 38:963–974.

Lakens D (2013) Calculating and reporting effect sizes to facilitate cumulative science: a practical primer for t-tests and ANOVAs. Front Psychol 4:863.

Liu D, Todorov E (2007) Evidence for the flexible sensorimotor strategies predicted by optimal feedback control. Journal of Neuroscience 27:9354–9368.

Milner TE, Franklin DW (2005) Impedance control and internal model use during the initial stage of adaptation to novel dynamics in humans. J Physiol 567:651–664.

Nashed JY, Crevecoeur F, Scott SH (2012) Influence of the behavioral goal and environmental obstacles on rapid feedback responses. J Neurophysiol 108:999–1009.

Nguyen KP, Zhou W, McKenna E, Colucci-Chang K, Bray LCJ, Hosseini EA, Alhussein L, Rezazad M, Joiner WM (2019) The 24-h savings of adaptation to novel movement dynamics initially reflects the recall of previous performance. J Neurophysiol 122:933–946.

Omrani M, Murnaghan CD, Pruszynski JA, Scott SH (2016) Distributed task-specific processing of somatosensory feedback for voluntary motor control. Elife 5.

Osu R, Hirai S, Yoshioka T, Kawato M (2004) Random presentation enables subjects to adapt to two opposing forces on the hand. Nat Neurosci 7:111–112.

Overduin SA, Richardson AG, Lane CE, Bizzi E, Press DZ (2006) Intermittent practice facilitates stable motor memories. Journal of Neuroscience 26:11888–11892.

Perich MG, Gallego JA, Miller LE (2018) A Neural Population Mechanism for Rapid Learning. Neuron 100:964–976 e967.

Pruszynski JA, Kurtzer I, Nashed JY, Omrani M, Brouwer B, Scott SH (2011) Primary motor cortex underlies multi-joint integration for fast feedback control. Nature 478:387–390.

Raymond JL, Medina JF (2018) Computational Principles of Supervised Learning in the Cerebellum. Annu Rev Neurosci 41:233–253.

Scheidt RA, Dingwell JB, Mussa-Ivaldi FA (2001) Learning to move amid uncertainty. J Neurophysiol 86:971–985.

Scott SH (2016) A Functional Taxonomy of Bottom-Up Sensory Feedback Processing for Motor Actions. Trends Neurosci 39:512–526.

Shadmehr R, Mussa-Ivaldi FA (1994) Adaptive representation of dynamics during learning of a motor task. Journal of Neuroscience 14:3208–3224.

Shadmehr R, Brashers-Krug T (1997) Functional stages in the formation of human long-term motor memory. J Neurosci 17:409–419.

Shadmehr R, Smith MA, Krakauer JW (2010) Error Correction, Sensory Prediction, and Adaptation in Motor Control. Annual Review of Neuroscience, Vol 33 33:89–108

Sheahan HR, Franklin DW, Wolpert DM (2016) Motor Planning, Not Execution, Separates Motor Memories. Neuron 92:773–779.

Singh K, Scott SH (2003) A motor learning strategy reflects neural circuitry for limb control. Nature Neuroscience 6:399–403.

Smith MA, Ghazizadeh A, Shadmehr R (2006) Interacting adaptive processes with different timescales underlie short-term motor learning. Plos Biology 4:1035–1043.

Stavisky SD, Kao JC, Ryu SI, Shenoy KV (2017) Motor Cortical Visuomotor Feedback Activity Is Initially Isolated from Downstream Targets in Output-Null Neural State Space Dimensions. Neuron 95:195–208 e199.

Thoroughman KA, Shadmehr R (2000) Learning of action through adaptive combination of motor primitives. Nature 407:742–747.

Todorov E (2005) Stochastic optimal control and estimation methods adapted to the noise characteristics of the sensorimotor system. Neural Computation 17:1084–1108.

Todorov E, Jordan MI (2002) Optimal feedback control as a theory of motor coordination. Nat Neurosci 5:1226–1235.

Wada Y, Kawabata Y, Kotosaka S, Yamamoto K, Kitazawa S, Kawato M (2003) Acquisition and contextual switching of multiple internal models for different viscous force fields. Neurosci Res 46:319–331.

Wagner MJ, Smith MA (2008) Shared Internal Models for Feedforward and Feedback Control. Journal of Neuroscience 28:10663–10673.

Wei K, Koerding K (2010) Uncertainty of feedback and state estimation determines the speed of motor adaptation. Frontiers in Computational Neuroscience 4.

Wolpert DM, Diedrichsen J, Flanagan JR (2011) Principles of sensorimotor learning. Nature Reviews Neuroscience 12:739–751.

